# String-Pulling as a Behavioral Assessment of Skilled Forelimb Motor Function in a Middle Cerebral Artery Occlusion Rat Model

**DOI:** 10.1101/2022.03.31.486586

**Authors:** Muriel Hart, Ashley A. Blackwell, Ian Q. Whishaw, Douglas G. Wallace, Joseph L. Cheatwood

**Affiliations:** Department of Anatomy, Southern Illinois University School of Medicine, Carbondale, Illinois, USA; Department of Psychology, Northern Illinois University, De Kalb, Illinois, 60115 USA; Canadian Centre for Behavioural Neuroscience, University of Lethbridge, Lethbridge, Alberta, Canada

**Keywords:** String-pulling, kinematics, posture, bilateral movement, MCAO, fine motor control, compensation

## Abstract

Stroke is a leading cause of long-term disability in humans and frequently results in bilateral impairments in fine motor control. Many behavioral tasks used to assess rodent models of stroke evaluate a single limb; however, recent work has demonstrated that bilateral hand-over-hand movements used to pull in a string assess skilled movement of both hands.

Devascularization focused on the forelimb portion of sensorimotor cortex has been observed to produce persistent disruptions in the topographical organization of string-pulling behavior. The current study examined changes in string-pulling after a more clinically relevant rodent model of stroke via middle cerebral artery occlusion (MCAO). Detailed movement analyses revealed disruptions in the bilateral organization of string-pulling and fine motor control of both hands. Rats missed the string more often with both hands, and when the string was missed on the impaired side, rats continued to cycle through subcomponents of string-pulling behavior as if the string were grasped in the hand. Rats also failed to make a grasping motion with the impaired hand when the string was missed and instead, demonstrated an open-handed raking-like motion. No differences were found in time to approach or to complete the string-pulling task to obtain a reward, demonstrating the importance of using a detailed functional analysis of movement to detect changes in performance. String-pulling behavior is sensitive at detecting changes in bilateral rhythmical hand control following MCAO providing a foundation for future work to investigate other models of stroke and to evaluate the efficacy of therapeutic interventions that enhance neuroplasticity.

## 1. Introduction

Stroke is a leading cause of long-term disability most commonly effecting the upper limbs acutely in 80% of patients and chronically in 40% of patients (Cramer et al., 1997). Disruptions in bimanual coordination and fine motor skills of the hands are reported by patients following stroke, to the extent that tasks involving the use of both hands are avoided (Sainburg et al., 2013; Lai et al., 2019). The most common strokes are ischemic and result in an occlusion of the middle cerebral artery (MCA). The MCA supplies blood to cortical and subcortical brain regions, including primary motor and somatosensory cortical areas responsible for the use of the face, trunk, and upper limbs (Navarro-Orozco et al., 2021). Middle cerebral artery occlusion (MCAO) in rodents is a common model to examine upper limb dysfunction, recovery, and therapeutics (Bederson et al., 1986; Kleim et al., 2007; Alaverdashvili, Whishaw, 2008; Schaar et al., 2010; Trueman et al., 2017). The ability to characterize performance following MCAO depends on several factors, including lesion extent/location and the type of task used to assess fine motor control (Gonzalez, Kolb, 2003; El Amki et al., 2017). While a variety of techniques currently exist to assess voluntary fine motor control in rodent models of stroke, most tasks fail to examine the concomitant use of both hands (Gonzalez et al., 2004). Thus, an examination of a bimanual task would model behavior that people engage in daily and provide a translational assessment of function and recovery.

Rodents spontaneously engage in bimanually coordinated string-pulling behavior that involves hand-over-hand movements to pull in a string to retrieve a food reward (Blackwell et al., 2018ab). String-pulling behavior provides a robust assessment of fine motor control, including functional performance measures and (i.e., task time, contacts, misses) and details about movement organization related to distance and direction, that is sensitive to focal cortical damage in rats (Blackwell et al., 2018c) and simulated space radiation exposure (Blackwell et al., 2021). For example, following forelimb sensorimotor devascularization, rats exhibit increased misses with the impaired hand and persistent deficits in distance and direction of movement. This focal unilateral damage shows some recovery with time and experience. The effects of the rodent MCAO model on bimanually coordinated movements remain to be determined and so is examined here. The objective of the study was to determine if string-pulling behavior is sensitive at detecting changes in movement organization with resulting recovery and compensation after MCAO.

For the study, rat string-pulling behavior was investigated after MCAO. Measures were made of the coordinated multiple upward reaches to grasp the string and downward withdraws to advance it. Reach and withdraw phases of string-pulling were decomposed into subcomponents (i.e., Advance, Grasp, Pull, Push and Lift), hand and mouth attempts to grasp the string were characterized as grasping, contacts or misses, and kinematic and topographic characteristics of movement were described as measures of the extent of deficits.

## 2. Methods

### 2.1. Subjects

A total of 15 adult male Long-Evans rats (Rattus norvegicus) began the study; rats that did not perform during the training sessions were categorized as non-pullers and excluded from the study (n = 3). Vivarium temperatures (20 to 21 C) and light (12-hr light-dark cycle) conditions were consistent throughout testing. Rats were food deprived for two nights prior to beginning string-pulling behavior and provided water ad libitum. All experimental protocols were approved by SIU Carbondale Institutional Animal Care and Use Committee.

### 2.2. Surgery

The middle cerebral artery occlusion surgery was performed using methods described previously (Cheatwood et al., 2011). Briefly, rats were anesthetized with isoflurane (5% in oxygen) and then placed in a stereotaxic device with an integrated anesthesia port. Rats were maintained on isoflurane (1% to 2.5% in oxygen) for the duration of the procedure. The left skull was exposed, and a craniotomy was made to allow access to the middle cerebral artery (MCA) close to where it exited the rhinal fissure. The MCA was then permanently ligated with a 10-0 suture and transected. Each animal then underwent a permanent occlusion of the common carotid artery (CCA) on the same side as the MCAO and a temporary occlusion (15 min.) of the contralateral CCA. Rats’ left hemispheres were chosen to receive MCAOs since patients more commonly experience strokes on the left side of the brain (Ma et al., 2020).

Similarly, patients with strokes more often present with permanent cerebral ischemia rather than reperfusion providing further basis for the model used in the current study. After the procedure, rats were removed from isoflurane anesthesia and allowed to recover before they were returned to their home cages.

### 2.3. Apparatus

The string-pulling apparatus was a transparent, rectangular box (19 cm x 19 cm). The apparatus sat on a table in a room with many cues. The rat remained in the testing apparatus for the entire string-pulling session within each day. In between each rat, the testing apparatus was thoroughly cleaned and prepared for the next rat.

### 2.4. Procedures

Rats were habituated to 0.5 m strings with unsalted cashews tied to the end draped in their home cage for two nights. Then, rats were trained on string-pulling across four consecutive days. String-pulling training sessions consisted of each rat pulling five 1.0 m strings within a 20-minute testing period. The following day, baseline performance was measured across four trials within a 20-minute testing session prior to surgeries. Rats were tested under the same conditions in string-pulling behavior on days 3, 7, and 14 following MCAO.

### 2.5. Behavioral Analysis

#### 2.5.1. General performance measures

Approach and pull times were used as general measures of performance in the string-pulling task. The amount of time it takes rats to approach the string once placed in the apparatus, and the amount of time it takes rats to pull in the string to reach the food item at the end are measures of motivation to engage in and complete the string-pulling task respectively.

#### 2.5.2. Hand movement analysis

String-pulling behavior is composed of phases of upward reaches away from the body to grasp the string and downward withdraws toward the body to pull in the string. String-pulling behavior is dependent on the ability of the hands to grasp the string to pull it in. The number of contacts (closing of digits around the string) was evaluated across testing. Misses that occurred when the rat failed to contact the string during the transition from the reach to withdraw phase was also measured across testing.

#### 2.5.3. Sequential movement analysis

A string-pulling cycle may be further decomposed into five different movement components: Advance, Grasp, Pull, Push, and Lift (see Figure 1). Rodents typically distribute movements evenly across all five components. The number of times rats engaged in each of the different movement components was recorded when contact was made with the string and divided by the total number of movements when contact was made with the string to create a ratio. Movement components were also separately recorded when rats missed the string using a ratio to compare the total number of each component during movement without the string to the total number of movements observed without the string when a miss occurred.

**Figure 1:**
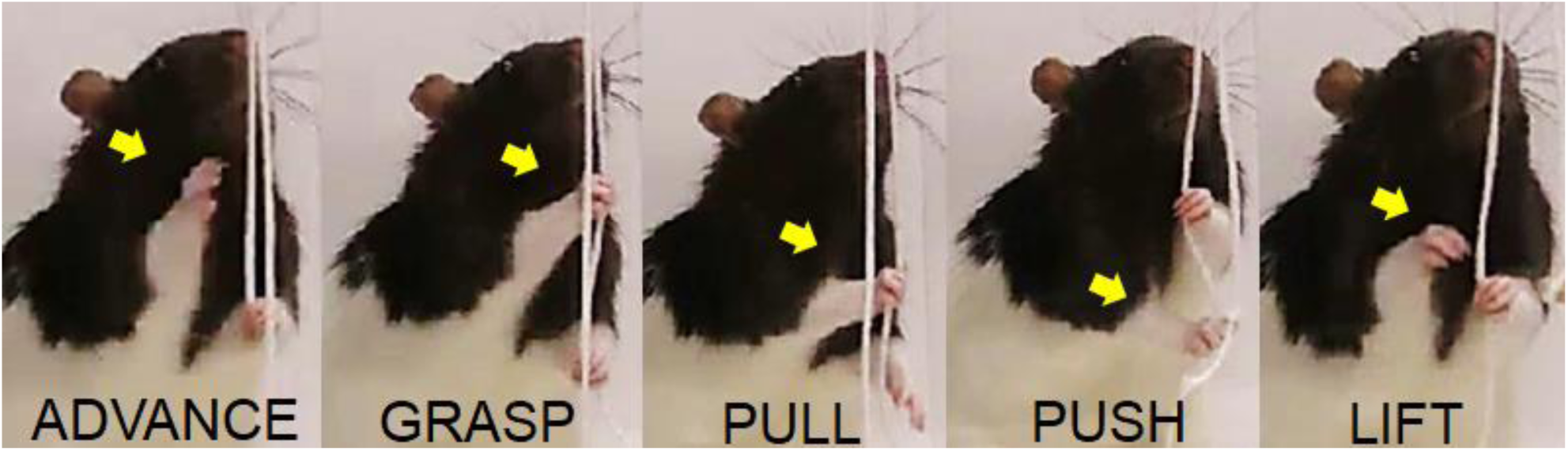
A right-hand Advance (A), Grasp (B), Pull (C), Push (D), and Lift (E) during movement with the string is displayed for a rat. All rats engaged in these subcomponents of movement.

#### 2.5.4. Motion capture analysis

String-pulling behavior was captured at 30 frames per second with a high-definition video camera (Canon Vixia HF21) for offline analysis. An open-source movement tracking software, Tracker (www.physlets.org), was used to manually digitize the left and right hands and the nose frame-by-frame during bouts of string-pulling behavior.

##### 2.5.4.1. Reach and withdraw component kinematic analysis

XY data generated by digitization via the Tracker program was segmented into reaches and withdraws based on the direction of movement (i.e., up or down). General (distance) and specific (path circuity, concentration, heading) measures of movement organization were quantified for reaches and withdraws with the left and right hands separately across testing. Distance traveled during reaches and withdraws was evaluated for the left and right hand for each of the sampled trials. Path circuity of reach and withdraw phases of movement was calculated by dividing the Euclidean distance by the total distance traveled. More direct paths yield values closer to 1.0 and more circuitous paths are closer to 0.0.

Next, circular statistics was used to evaluate the concentration of reach and withdraw endpoints and their heading directions. The variability of the directional heading of reach and withdraw phases of movement were calculated using parameter of concentration. Circular statistics were used to quantify the consistency of heading directions for sampled string-pulling trials within each testing day (Batschelet, 1981). The start and end coordinates for both phases of movement were transformed such that the start of the path was the origin (0,0), and the angle of the end coordinate was calculated relative to a polar coordinate system (0: right; 90: up; 180: left; 270: down). Values range from 0.0 (variable heading that are uniformly distributed all directions) to 1.0 (no variability in heading). These values were used to calculate average parameter of concentration for each day across testing.

Reach and withdraw phases of movement are oriented in a specific direction. The directional heading of movement was evaluated by transforming the start and end coordinates of the path such that the start of the path is the origin (0,0), and the angle of the end coordinate is calculated relative to a polar coordinate system (0: right; 90: up; 180: left; 270: down).

##### 2.5.4.2. Nose kinematic analysis

The nose was also tracked using DLC. Then, XY data exported from DLC was used to evaluate nose kinematics between groups. The maximum and minimum X and Y range of movement with the nose was quantified across bouts of string-pulling behavior for each day of testing.

### 2.6. Histology

Following behavioral testing, rats were deeply anesthetized and perfused with phosphate-buffered saline, followed by 4% paraformaldehyde. Brains were stored in paraformaldehyde solution for 24 hr and then moved to a 30% sucrose solution for approximately 48 hr. Then, brains were sliced in to 40 μm sections using a vibratome and stained with cresyl violet to investigate lesion volume.

### 2.7. Statistical analysis

All statistical analyses included the repeated measures of day that reflected post-surgery days 3, 7, and 14 of testing. Sequential movement analyses for MCAO rats included the repeated measure Hands, designated as left and right. Repeated measures ANOVAs were used to evaluate main effects and interactions on each dependent measure in the string-pulling. The Greenhouse-Geisser (G-G) correction was used in analyses where Mauchly’s test indicated significant departure from the assumption of sphericity. Partial eta squared (η^2^p) was used as a measure of effect size for each main effect and interaction. Linear trend and Tukey HSD tests were used for post-hoc analyses.

## 3. Results

### 3.1. Histology

Representative photographs depict minimum (red) and maximum (black) lesion extent following MCAO (see Figure 2A). Lesion volume was calculated for each rat that received a MCAO (see Figure 2B). Extent of lesions ranged from ∼16.8% to 30.95%. Mainly cortical regions were affected by MCAO, including primary and secondary sensorimotor areas important for skilled hand use (Edwards et al., 2019). Additional cortical brain regions damaged by MCAO encompassed of the barrel cortex responsible for vibrissae movement and involved in sensorimotor processing (Peterson, 2019).

**Figure 2:**
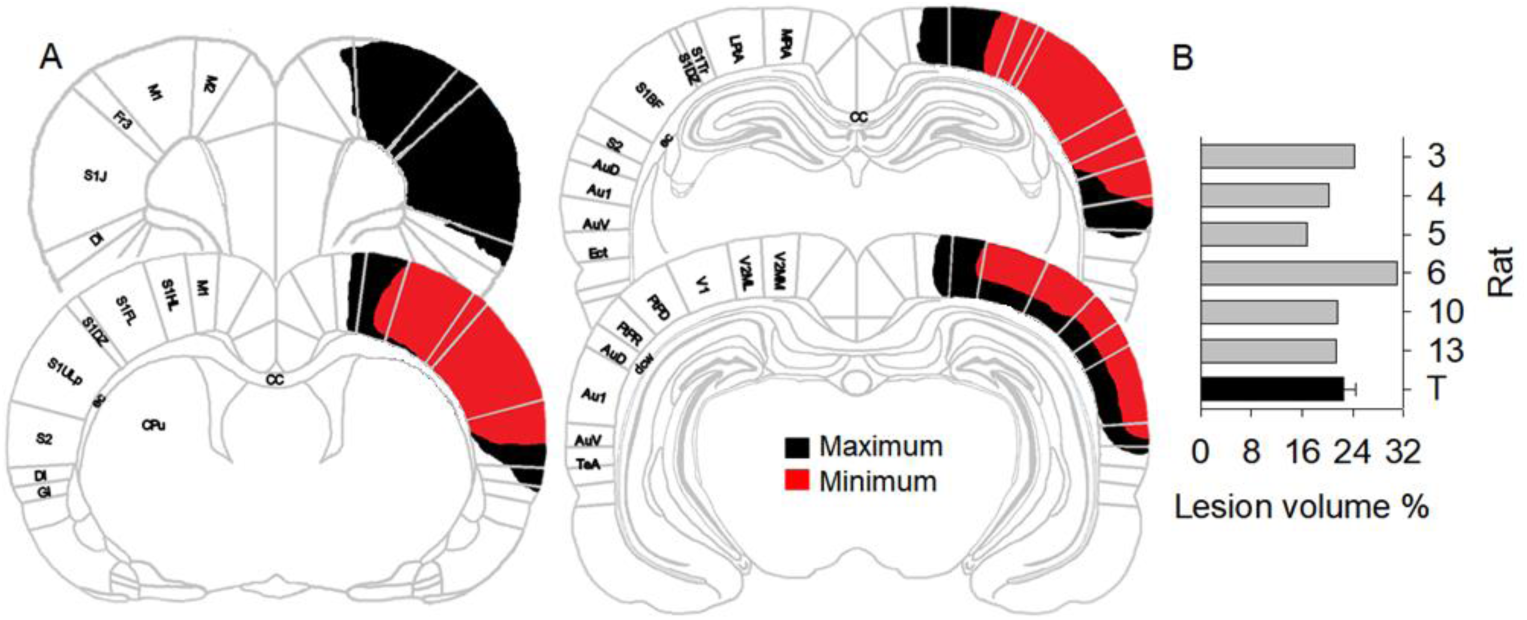
Minimum (red) and maximum (black) lesion extent is shown (A) with lesion volume as a percentage for each rat that received a MCAO (B).

### 3.2. String-pulling general behavioral analysis

Independent samples t-tests were used to compare performance between the sham and MCAO group during baseline testing, prior to MCAO, on each measure. No differences were found between groups on any measures during baseline; therefore, repeated measures ANOVAs were used to evaluate performance between groups during post-tests on days 3, 7 and 14.

Broad motivational and endpoint measures of performance were characterized to evaluate group differences in the organization of bimanual string-pulling behavior across (see Table 1). Approach time and pull duration were evaluated across testing between groups as broad motivational measures of performance. The G-G correction (ε=0.513) was used to adjust the degrees of freedom associated with the lack of sphericity in pull duration. Separate Repeated-measures ANOVAs conducted on approach time and pull duration failed to reveal significant main effects of Group, Day, or Group by Day interactions. Rats took similar amounts of time to approach the string once placed in the testing apparatus as well as similar pull durations across testing

**Table 1.** Motivational analysis and endpoint measures (*p < 0.05).

| Time | Approach | $f$ | $df$ | $p$ | $\eta^2p$ |
| --- | --- | --- | --- | --- | --- |
|  | Day | 2.323 | 2, 20 | 0.124 | 0.189 |
|  | Day X Group | 0.279 | 2, 20 | 0.760 | 0.027 |
|  | Group | 0.341 | 1, 10 | 0.572 | 0.033 |
|  | Pull |  |  |  |  |
|  | Day | 2.140 | 2, 20 | 0.174 | 0.176 |
|  | Day X Group | 1.810 | 2, 20 | 0.208 | 0.153 |
|  | Group | 2.182 | 1, 10 | 0.170 | 0.179 |
| Contacts | Left-hand |  |  |  |  |
|  | Day | 2.483 | 2, 20 | 0.109 | 0.199 |
|  | Day X Group | 0.784 | 2, 20 | 0.470 | 0.073 |
|  | Group | 0.206 | 1, 10 | 0.660 | 0.020 |
|  | Right-hand |  |  |  |  |
|  | Day | 2.163 | 2, 20 | 0.141 | 0.178 |
|  | Day X Group | 1.484 | 2, 20 | 0.251 | 0.129 |
|  | Group | 3.804 | 1, 10 | 0.080 | 0.276 |
| Misses | Left-hand |  |  |  |  |
|  | Day | 2.136 | 2, 20 | 0.144 | 0.176 |
|  | Day X Group | 1.009 | 2, 20 | 0.382 | 0.092 |
|  | Group | 24.258 | 1, 10 | *0.001 | 0.708 |
|  | Right hand |  |  |  |  |
|  | Day | 1.380 | 2, 20 | 0.275 | 0.121 |
|  | Day X Group | 1.537 | 2, 20 | 0.239 | 0.133 |
|  | Group | 53.376 | 1, 10 | *<0.001 | 1.000 |

Rats in both groups exhibited misses while attempting to contact and grasp the string to pull it in with the left (see Figure 3A) and right hand (see Figure 3B). The number of misses made with the left-and right-hands while pulling in the string were evaluated across testing between groups. Separate Repeated-measures ANOVAs conducted on left- [F (1, 10) = 24.258, p = 0.001, η^2^p = 0.708] (see Figure 3C) and right-hand [F (1, 10) = 53.376, p = 0.001, η^2^p = 1.000] (see Figure 3D) misses revealed significant main effects of Group yet failed to reveal significant main effects of Day or Group by Day interactions. Following MCAO, rats exhibited an increase in misses, or failures to contact the string, with both the left- and right-hands compared to sham rats across testing.

**Figure 3:**
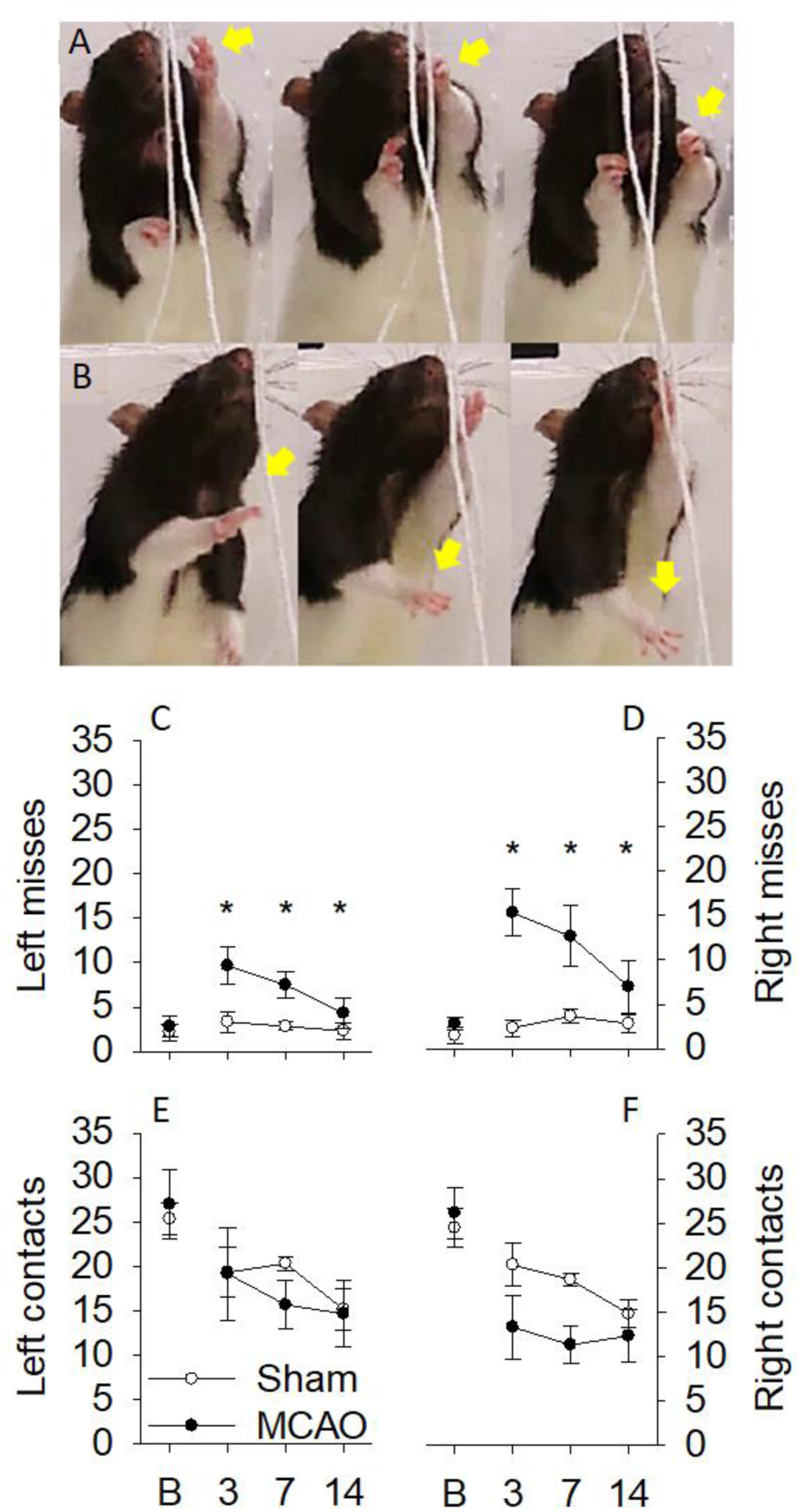
A frame-by-frame representative left-(A) and right-hand (B) miss is displayed; notice that the hand and fingers grasp even when the string is missed (A) or exhibit extensor spasticity (B) across subcomponents of movement (i.e., Grasp, Pull, Push). Following MCAO, rats engaged in significantly more misses with the left (C) and right hands (D) relative to sham rats across testing, while left-(E) and right-hand (F) contacts were similar between groups. *p < 0.050

### 3.3. Sequential movement analyses

#### 3.3.1. Contacts

Rats sequentially organize string-pulling behavior into subcomponents of movements, including an Advance to Grasp the string, a Pull and Push to move the string into the apparatus, and a release to Lift the hand and begin the next pulling cycle (see Figure 1). A sequential analysis of the left- and right-hands revealed all rats engaged in subcomponents of movement when contact was made with the string with two exceptions (see Table 2). Separate repeated measures ANOVAs revealed that the number of pushes to further advance the string into the apparatus during withdraw movements significantly differed between Groups for the left [F (2, 20) = 7.670, p = 0.020, η^2^ _p_ = 0.434] and right [F (2, 20) = 10.559, p = 0.009, η^2^ _p_ = 0.514] hands (see Table 2).

**Table 2.**
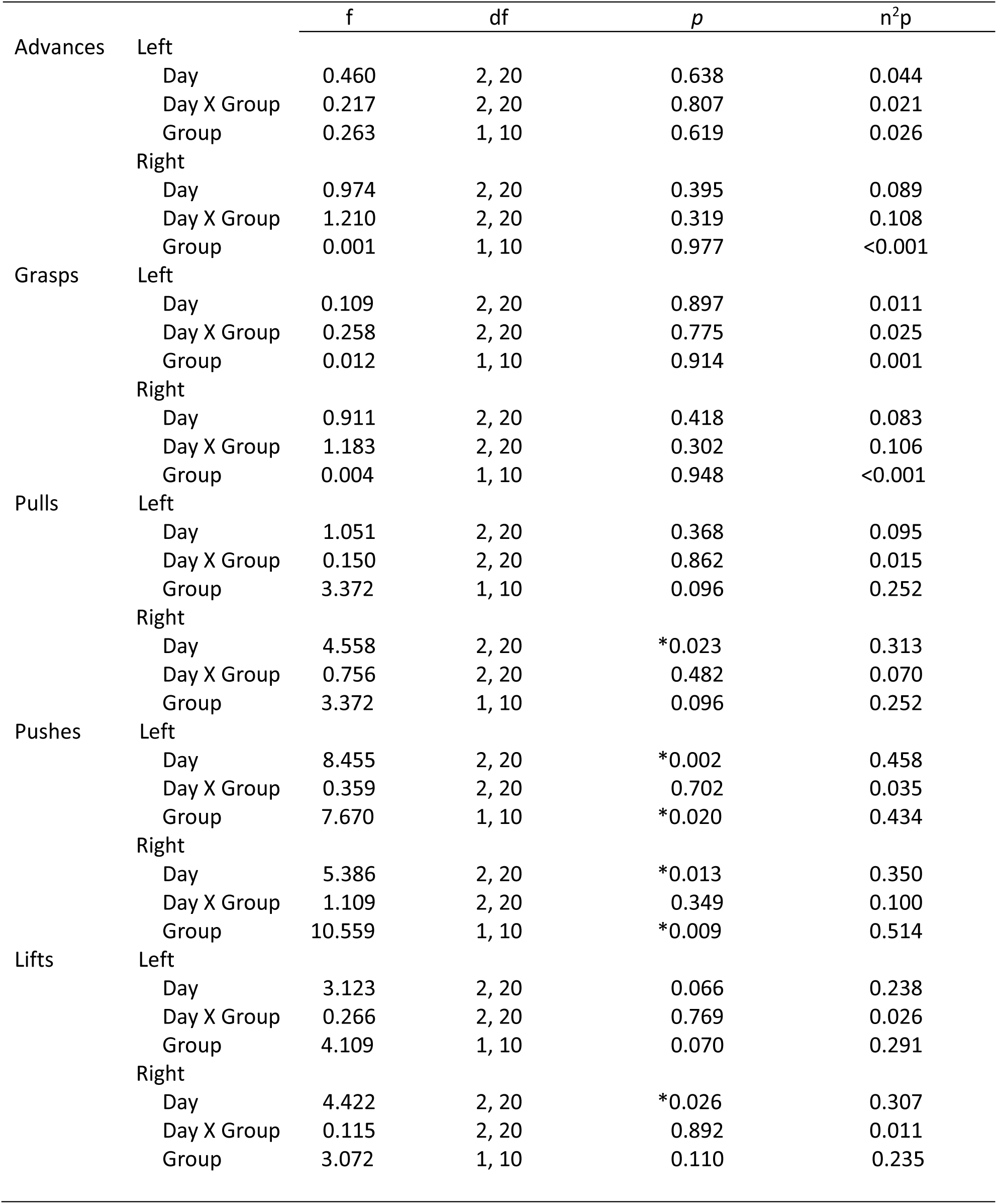
Sequential movement analysis between groups (*p < 0.05).

#### 3.3.2. Misses

Next, separate movement analyses conducted when MCAO rats missed the string revealed changes in subcomponents between the left and right hands across testing (see Table 3). Rats in the sham group did not engage in enough misses to investigate subcomponents of movement organization. Therefore, sequential analyses of movement subcomponents during misses were only conducted for rats that received a MCAO to further investigate the nature of misses with the left- and right-hands.

**Table 3.**
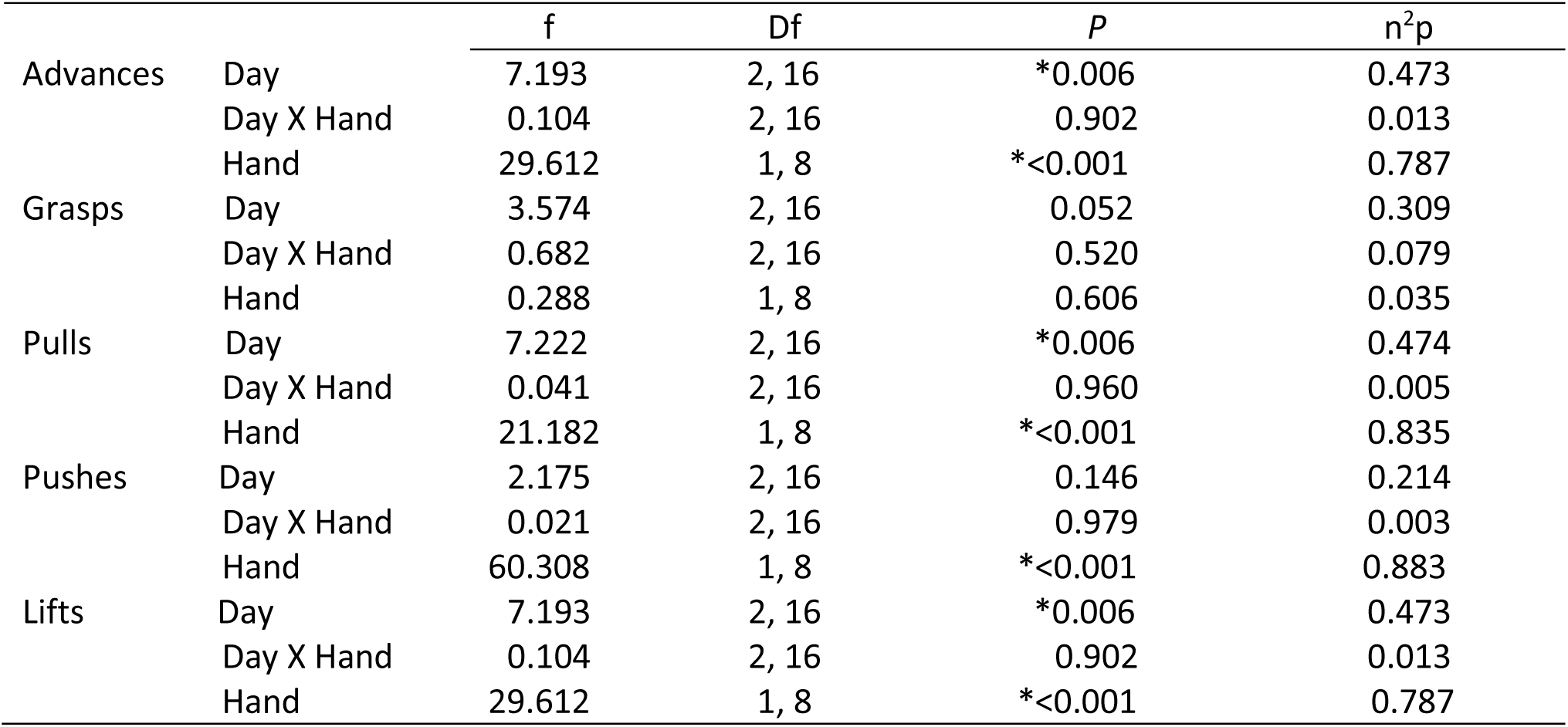
Sequential movement analysis during misses for rats that received MCAO (*p < 0.05).

##### 3.3.2.1. Advances

Advances toward the string with the left- and right-hands were evaluated during misses for rats that received a MCAO across testing (see Figure 4A). A Repeated measures ANOVA conducted on advances revealed that MCAO rats engaged in more left- than right-hand advances across testing [F (1, 8) = 29.612, p < 0.001, η^2^_p_ = 0.787] (see Figure 4B). Further, left- and right-hand advances increased as a function of day [F (2, 16) = 7.193, p = 0.006, η^2^_p_ = 0.473], but there was not a significant Hand by Day interaction.

**Figure 4:**
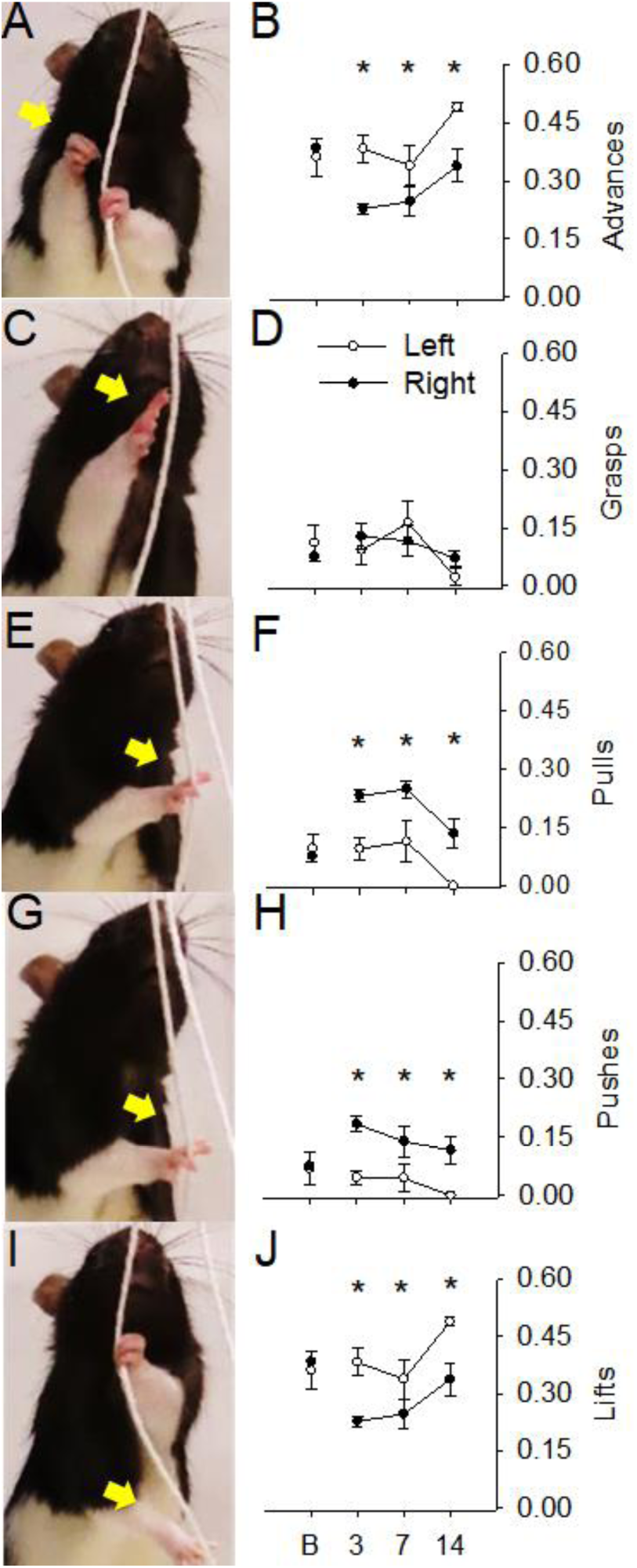
A right-hand Advance (A), Grasp (B), Pull (C), Push (D), and Lift (E) during movement without the string (i.e., during a miss with extensor spasticity) is displayed along with corresponding left and right-hand graphs for rats that received a MCAO. MCAO resulted in greater advances and lifts with the left-hand and less pulls and pushes relative to the right-hand. *p < 0.050

##### 3.3.2.2.Grasps

After MCAO, rats continued to engage in grasps even when the string was missed with the left-and right-hands (see Figure 4C). However, a Repeated measures ANOVA conducted on grasps failed to reveal a significant main effect of Day, Hand, or Day by Hand interaction (see Figure 4D).

##### 3.3.2.3. Pulls

When rats missed the string following MCAO they cycled through pulls with the left- and right-hand (see Figure 4E). A Repeated measures ANOVA conducted on pulls revealed a significant main effect of Hand [F (1, 8) = 21.182, p < 0.001, η^2^_p_ = 0.835] and Day [F (2, 16) = 7.222, p < 0.006, η^2^_p_ = 0.474] yet failed to reveal a significant Hand by Day interaction (see Figure 4F). Rats that received a MCAO engaged in more right-than left-hand pulls across testing. Overall, pulls with both hands decreased across testing.

##### 3.3.2.4. Pushes

Rats also continued to engage in pushes with the left- and right-hand during misses (see Figure 4G). A Repeated measures ANOVA conducted on pushes revealed a significant main effect of Hand [F (1, 8) = 60.308, p < 0.001, η^2^_p_ = 0.883] yet failed to reveal a significant main effect of Day or Hand by Day interaction (see Figure 4H). Following MCAO, rats engaged in more right-than left-hand pushes across testing.

##### 3.3.2.5. Lifts

Lifts with the left- and right-hand during misses were also evaluated across testing (see Figure 4I). A Repeated measures ANOVA conducted on lifts revealed a significant main effect of Hand and Day yet failed to reveal a significant Hand by Day interaction (see Figure 4J). Rats that received a MCAO engaged in more right-than left-hand lifts across testing. Further, lifts with both hands decreased as a function of day.

### 3.4. Motion capture analysis

#### 3.4.1. Nose kinematic analysis

While MCAO rats engaged in subcomponents of movement (Advance, Grasp, Pull, Push, and Lift) during bouts of string-pulling behavior, the organization of these movements were drastically altered. Importantly, following MCAO, rats engaged in whole-body movements to initiate and perform all subcomponents of movement (see Figure 5).

**Figure 5:**
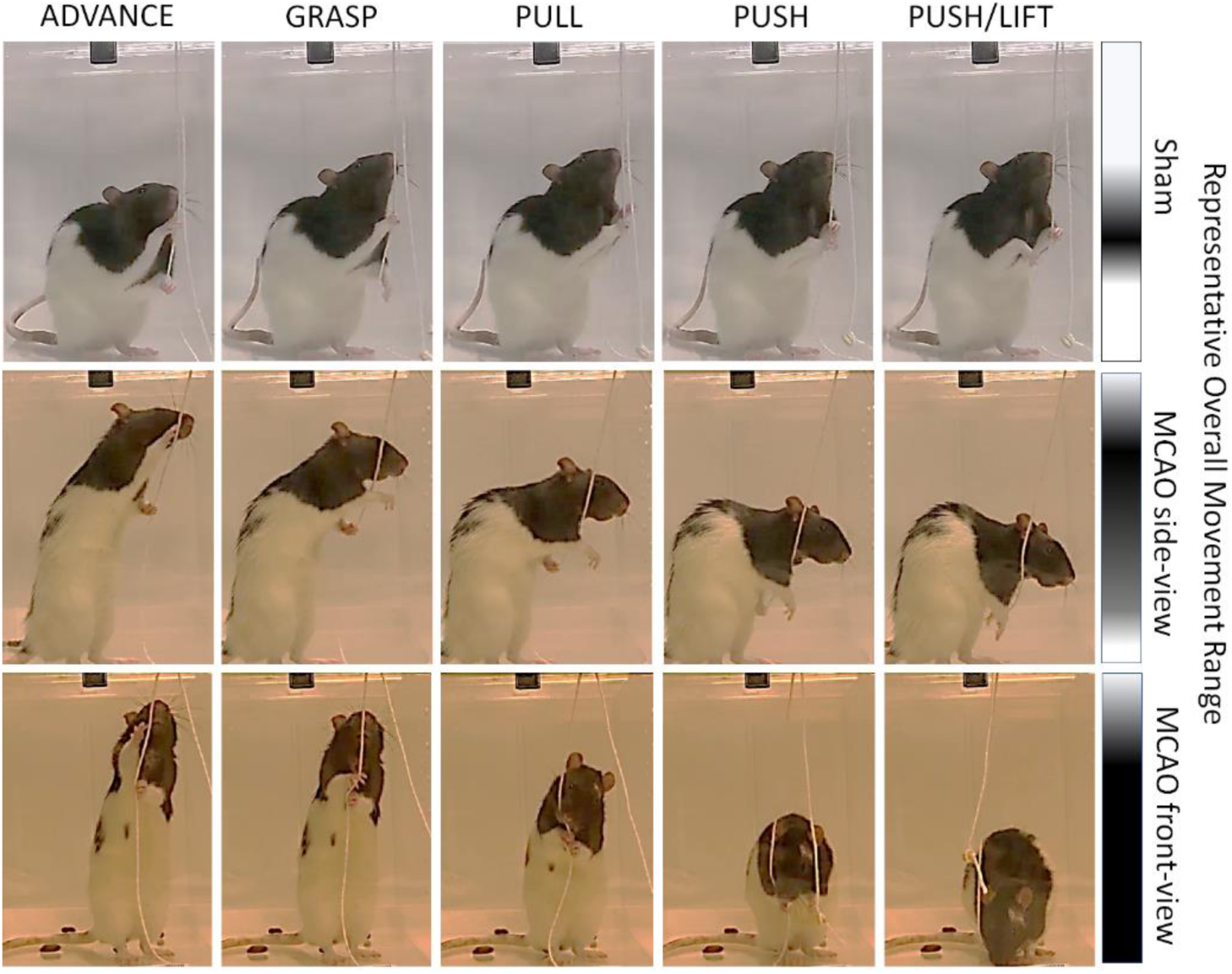
Overall range in movement is demonstrated across subcomponents of movement (Advance, Grasp, Pull, Push, Lift) for a representative sham rat (top row) and two rats that received MCAO (middle: side view and bottom row: front view). The bars to the right of each row show the range of y-axis movement for a bout of string-pulling behavior for each representative rat. Black represents areas where most of the movement occurred, while the gray areas denote less movements, and white signifies no movements were made within that region.

Whether the string was contacted or missed with the hands, rats that received a MCAO used their entire body to advance the string. Typically, sham rats occasionally use their mouths to contact the string to pull it in and sometimes (more rarely) use their entire body to aid in advancing the string. While no significant differences were observed in mouth contacts in the current study (see Table 1), after MCAO, rats (see Figure 6A) moved their noses differently relative to sham rats (see Figure 6B).

**Figure 6:**
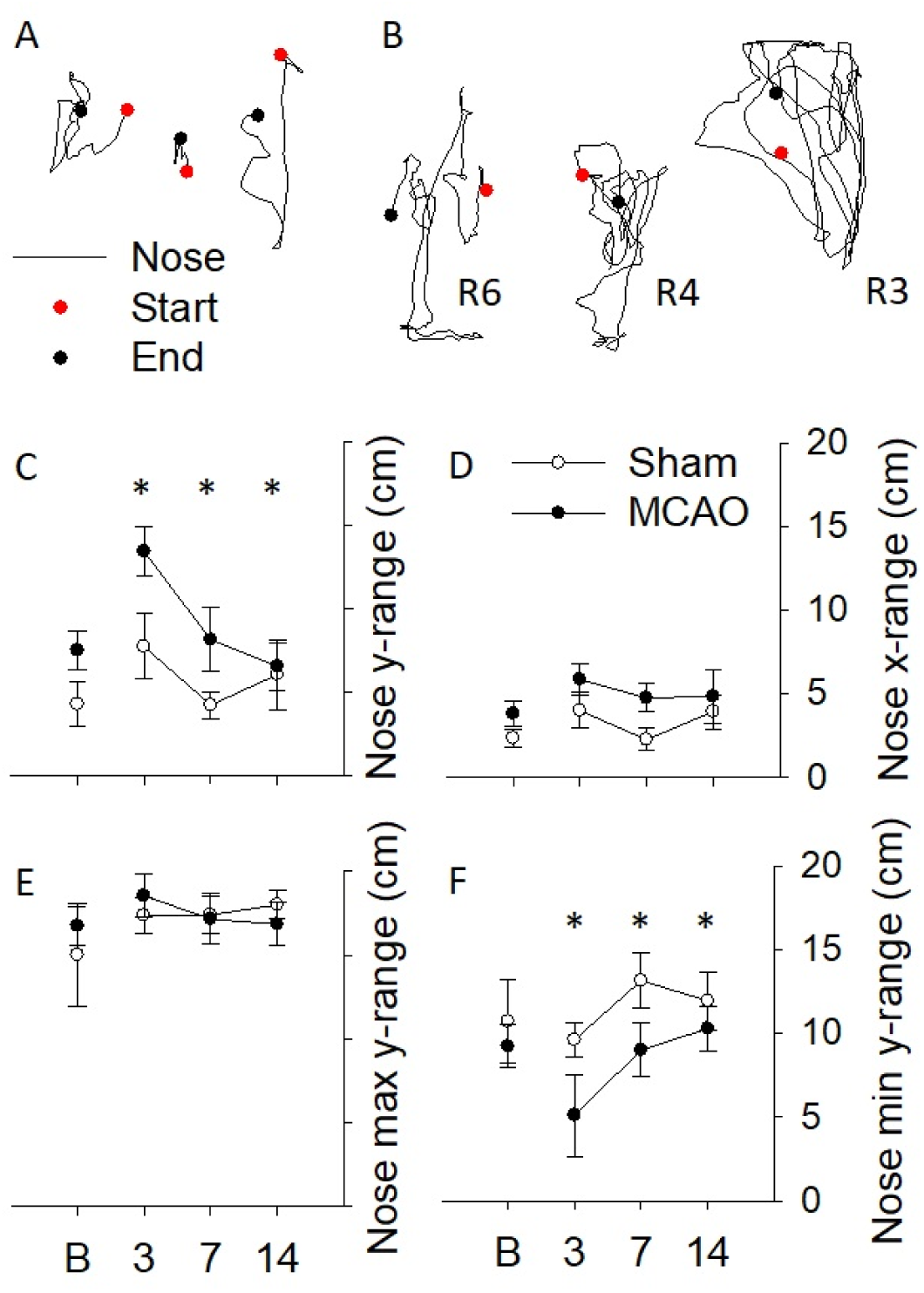
Topography of the nose is plotted from a bout of string-pulling behavior for representative sham (A) and MCAO (B) rats. Movement of the nose in the y-range (C) significantly differed between groups with rats traveling further distances after MCAO, while no changes were present in the x-range of nose movement. Further investigation of the min and max range of nose movement within the y-axis revealed that after MCAO rats exhibited a significantly lower min y-range. *p < 0.050

A repeated measures ANOVA conducted on the y-range of nose movement revealed a significant effect of Group [F (1, 10) = 5.079, p = 0.048, η^2^p = 0.338] and Day [F (2, 20) = 5.004, p = 0.017, η^2^p = 0.333] yet failed to reveal a significant Group by Day interaction [F (2, 20) = 1.387, p = 0.273, η^2^p = 0.122] (see Figure 6C). Following MCAO, rats traveled further distances in y-range movement with the nose, and this distance decreased across testing for both groups. The minimum and maximum y-range movement of the nose was evaluated to further investigate the nature of this difference. A repeated measures ANOVA conducted on the minimum y-range of the nose revealed a significant effect of Group [F (1, 10) = 6.079, p = 0.033, η^2^p = 0.378] without a significant Group by Day interaction [F (2, 20) = 0.430, p = 0.567, η^2^p = 0.041] or effect of Day [F (2, 20) = 3.293, p = 0.058, η^2^p = 0.248] (see Figure 6F). MCAO rats traveled further distances in the minimum y-range moving all the way down to the floor at the bottom of the apparatus to advance the string, while sham rats remained upright to pull in the string.

In contrast, no differences were observed in the maximum y-range of the nose: Group [F (1, 10) = 0.004, p = 0.954, η^2^p = 3.541e-4], Day [F (2, 20) = 0.306, p = 0.740, η^2^p = 0.030], and Group by Day interaction [F (2, 20) = 0.822, p = 0.454, η^2^p = 0.076] suggesting rats reached to similar heights to manipulate the string (see Figure 6E). Similarly, a repeated measures ANOVA conducted on the x-range of nose movement failed to reveal any significant differences: Group [F (1, 10) = 2.380, p = 0.154, η^2^p = 0.192], Day [F (2, 20) = 1.382, p = 0.274, η^2^p = 0.121], Group by Day [F (2, 20) = 0.415, p = 0.666, η^2^p = 0.040] (see Figure 6D). Horizontal nose movement was similar between groups across testing.

##### 3.4.2. Reach component

All rats engaged in left- and right-hand reaches to grasp the string with MCAO rats exhibiting selective changes in movement organization during reach paths (see Table 4). Distance traveled during reaches (see Figure 7C-D) and the circuity of reach paths were similar between MCAO and sham rats for the left and right hands. The G-G correction method was used to adjust the degrees of freedom associated with the lack of sphericity in distance traveled (ε=0.626) with the left-hand and in path circuity with the right-hand (ε=0.596). Further, the endpoints of reaches were similarly concentrated between groups (see Figure 8).

**Table 4.** Reach kinematics (*p < 0.05).

| Distance |  | f | df | p | n<sup>2</sup>p |
| --- | --- | --- | --- | --- | --- |
| Distance | Left |  |  |  |  |
|  | Day | 3.213 | 1.253, 10.023 | 0.098 | 0.287 |
|  | Day X Group | 0.571 | 1.253, 10.023 | 0.505 | 0.067 |
|  | Group | <0.001 | 1, 8 | 0.983 | 0.313 |
|  | Right |  |  |  |  |
|  | Day | 0.640 | 2, 16 | 0.540 | 0.074 |
|  | Day X Group | 1.629 | 2, 16 | 0.227 | 0.169 |
|  | Group | 0.144 | 1, 8 | 0.714 | 0.018 |
| Path circuitry | Left |  |  |  |  |
|  | Day | 1.854 | 2, 16 | 0.189 | 0.188 |
|  | Day X Group | 0.297 | 2, 16 | 0.747 | 0.036 |
|  | Group | 0.741 | 1, 8 | 0.414 | 0.085 |
|  | Right |  |  |  |  |
|  | Day | 0.259 | 1.192, 9.534 | 0.663 | 0.031 |
|  | Day X Group | 0.819 | 1.192, 9.534 | 0.409 | 0.093 |
|  | Group | 0.027 | 1, 8 | 0.875 | 0.003 |
| Concentration | Left |  |  |  |  |
|  | Day | 0.482 | 2, 16 | 0.626 | 0.057 |
|  | Day X Group | 1.114 | 2, 16 | 0.352 | 0.122 |
|  | Group | 2.680 | 1, 8 | 0.140 | 0.251 |
|  | Right |  |  |  |  |
|  | Day | 1.124 | 2, 16 | 0.349 | 0.123 |
|  | Day X Group | 2.290 | 2, 16 | 0.133 | 0.223 |
|  | Group | 1.534 | 1, 8 | 0.251 | 0.161 |
| Heading | Left |  |  |  |  |
|  | Day | 1.593 | 2, 16 | 0.234 | 0.166 |
|  | Day X Group | 4.045 | 2, 16 | *0.038 | 0.336 |
|  | Group | 2.843 | 1, 8 | 0.130 | 0.262 |
|  | Right |  |  |  |  |
|  | Day | 0.145 | 2, 16 | 0.589 | 0.064 |
|  | Day X Group | 0.615 | 2, 16 | 0.553 | 0.170 |
|  | Group | 0.010 | 1, 8 | 0.912 | 0.114 |

**Figure 7:**
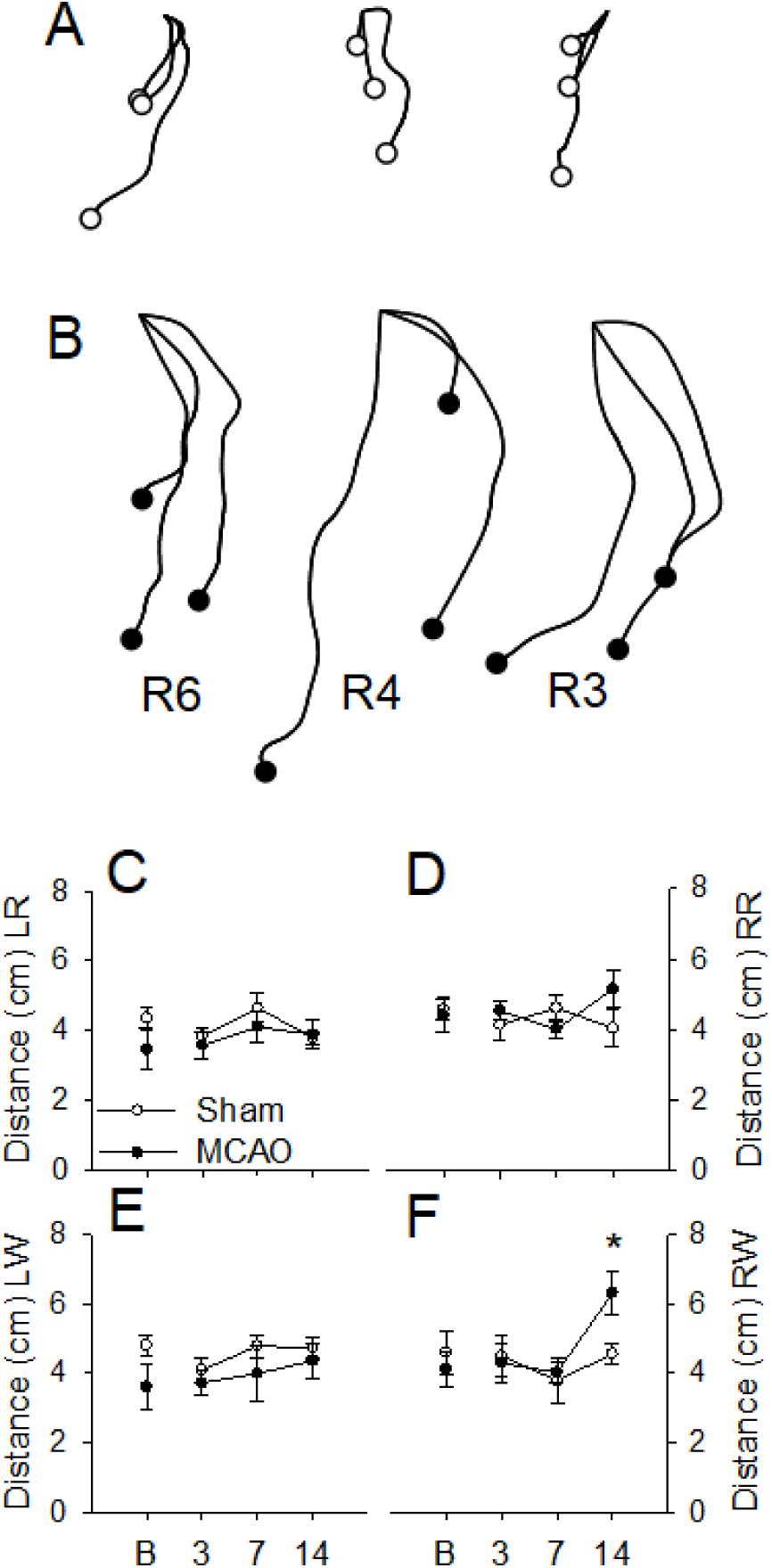
Representative topographic plots of the right-hand during withdraws is shown for sham (A) and MCAO (B) rats from day 14 of testing. Filled-in circles represent the endpoints of the withdraw paths. Distance traveled with the left- (LR; C) and right-hand during reaches (RR; D) as well as left- (LW; E) and right-hand during withdraws (RW; F) are graphed across testing. Following MCAO, rats traveled further distances during right-hand withdraws on day 14 of testing. *p < 0.050

**Figure 8:**
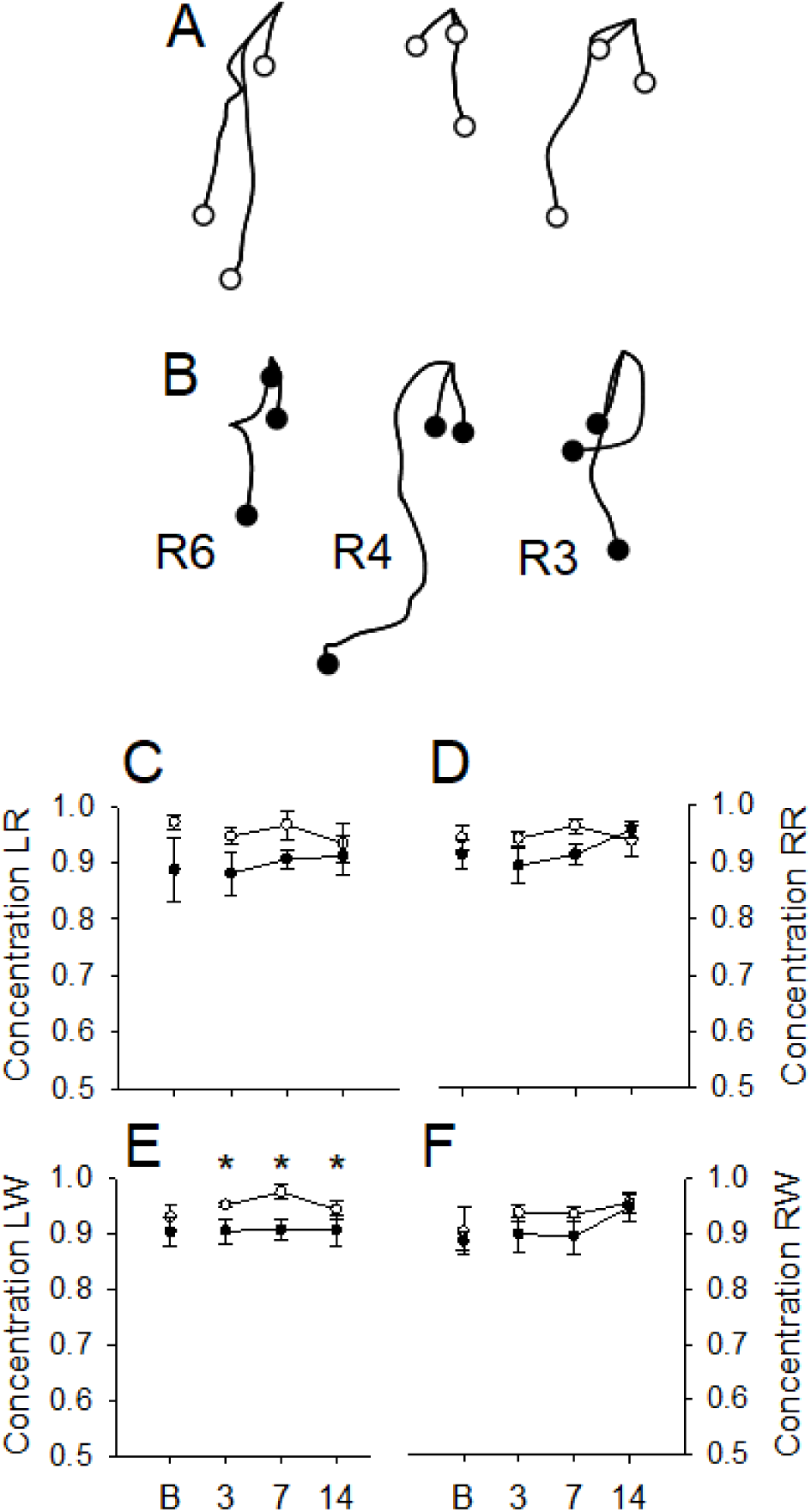
Representative topographic plots of left-hand withdraws is shown for sham (A) and MCAO (B) rats. Filled-in circles represent the endpoints of the withdraw paths. Concentration of movement with the left- (LR; C) and right-hand during reaches (RR; D) as well as left-(LW; E) and right-hand during withdraws (RW; F) are shown across testing. After MCAO, rats exhibited

Following MCAO, rats exhibited changes in heading direction of reaches that were specific to the left-hand (see Figure 9). A Repeated-measures ANOVA conducted on left-hand heading revealed a significant Group by Day interaction [F (2, 16) = 4.045, p = 0.038, η^2^p = 0.336] yet failed to reveal a significant main effect of Group or Day (see Table 4). Post hoc analyses revealed that rats in the MCAO group exhibited greater heading with the left-hand during the reach phase on days 3 and 7 of testing compared to rats in the sham group (HSD < 0.05); however, heading did not differ between groups on day 14 of testing. A Repeated-measures ANOVA conducted on right-hand heading failed to reveal any significant differences (see Table 4).

**Figure 9:**
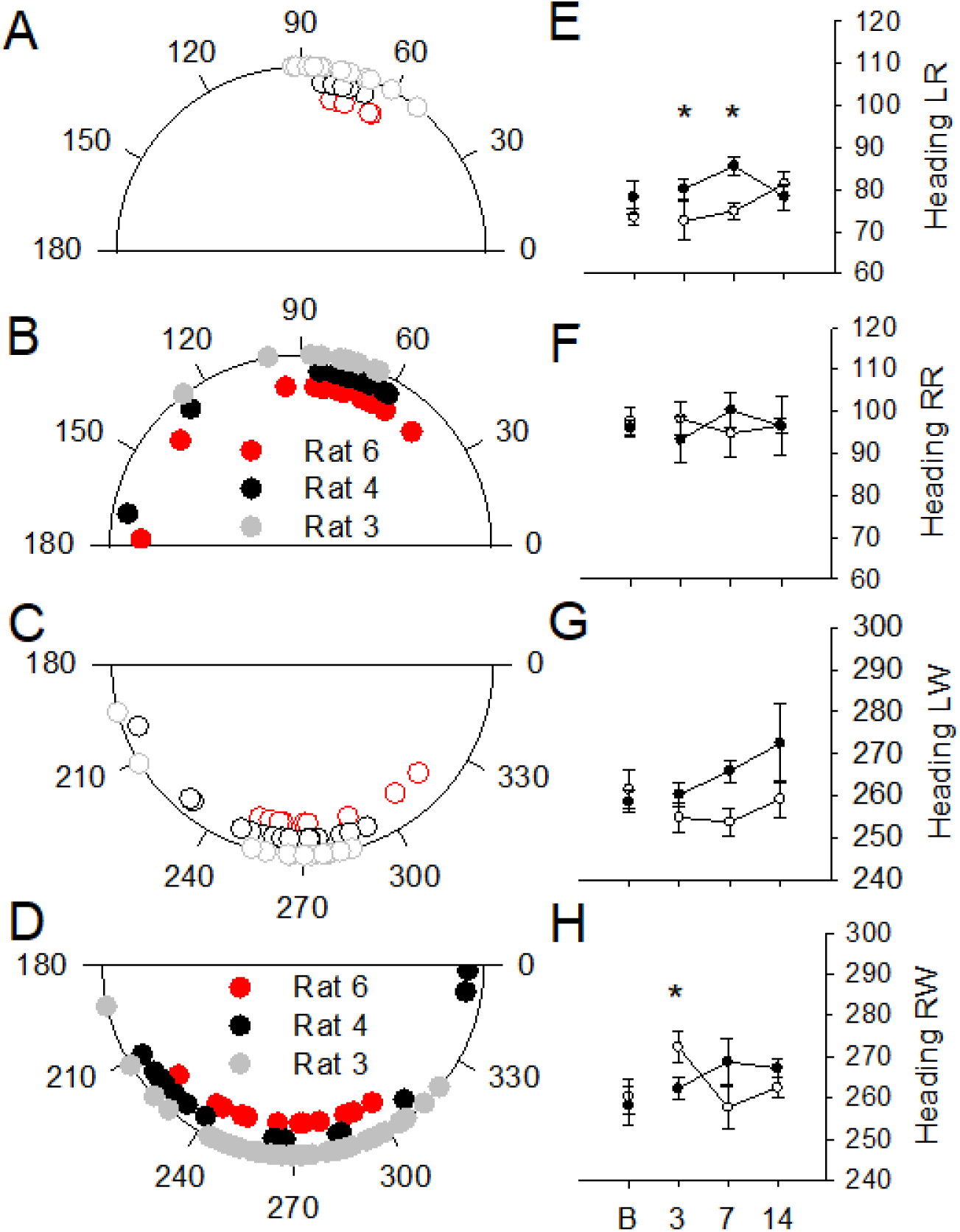
Representative circular plots of left-hand reaches are shown for sham (A) and MCAO (B) rats along with right-hand withdraws (C, D). Heading direction of movement with the left-(LR; E) and right-hand during reaches (RR; F) and the left- (LW; G) and right-hand during withdraws (RW; H) are shown across testing. Rats exhibited changes in L heading direction on days 3 and 7 after MCAO as well as RW heading on day 3 of testing. *p < 0.050

##### 3.4.3. Withdraw component

All rats engaged in withdraws with both hands to advance the string. Distance traveled with the left- and right-hands during the withdraw phase of movement were evaluated on day 3, 7, and 14 between groups (see Table 5). A Repeated-measures ANOVA conducted on left-hand distance revealed a significant main effect of Day [F (2, 16) = 4.735, p = 0.024, η^2^p = 0.372] yet failed to reveal a significant main effect of Group or Group by Day interaction. Post hoc analyses revealed a significant increasing linear trend in left-hand distance across testing [*F* (1, 8) = 11.503, *p* = 0.009, η^2^p =0.590]. Distance traveled increased with the left hand during the withdraw phase of movement for both groups across testing.

**Table 5.** Withdraw kinematics (*p < 0.05).

| Distance | Left | f | df | p | $\eta^2p$ |
| --- | --- | --- | --- | --- | --- |
|  | Day | 4.735 | 2, 16 | *0.024 | 0.372 |
|  | Day X Group | 0.054 | 2, 16 | 0.948 | 0.007 |
|  | Group | 0.326 | 1, 8 | 0.584 | 0.039 |
|  | Right |  |  |  |  |
|  | Day | 6.051 | 2, 16 | *0.011 | 0.431 |
|  | Day X Group | 3.741 | 2, 16 | *0.046 | 0.319 |
|  | Group | 0.500 | 1, 8 | 0.499 | 0.096 |
| Path circuitry | Left |  |  |  |  |
|  | Day | 0.187 | 2, 16 | 0.813 | 0.023 |
|  | Day X Group | 1.159 | 2, 16 | 0.339 | 0.127 |
|  | Group | 1.935 | 1, 8 | 0.202 | 0.195 |
|  | Right |  |  |  |  |
|  | Day | 4.814 | 2, 16 | *0.023 | 0.376 |
|  | Day X Group | 0.023 | 2, 16 | 0.122 | 0.232 |
|  | Group | 4.520 | 1, 8 | 0.066 | 0.361 |
| Concentration | Left |  |  |  |  |
|  | Day | 0.523 | 2, 16 | 0.603 | 0.061 |
|  | Day X Group | 0.981 | 2, 16 | 0.397 | 0.109 |
|  | Group | 7.813 | 1, 8 | *0.023 | 0.494 |
|  | Right |  |  |  |  |
|  | Day | 0.930 | 2, 16 | 0.415 | 0.104 |
|  | Day X Group | 0.629 | 2, 16 | 0.546 | 0.073 |
|  | Group | 2.508 | 1, 8 | 0.239 | 0.930 |
| Heading | Right |  |  |  |  |
|  | Day | 0.671 | 2, 16 | 0.525 | 0.077 |
|  | Day X Group | 4.299 | 2, 16 | *0.032 | 0.350 |
|  | Group | 0.767 | 1, 8 | 0.407 | 0.087 |

Next, a Repeated-measures ANOVA conducted on right-hand distance revealed a significant main effect of Day [F (2, 16) = 6.051, p = 0.011, η^2^p = 0.431] and Group by Day interaction [F (2, 16) = 3.741, p = 0.046, η^2^p = 0.319] yet failed to reveal a significant main effect of Group. Separate trend analyses were conducted on each group’s right-hand distance traveled to further investigate the nature of this interaction. Post hoc trend analysis revealed that right-hand distance traveled for rats in the MCAO group significantly increased across testing [*F* (1, 4) = 8.872, *p* = 0.041, η^2^p = 0.688] (see Figure 7A), while distance for rats in the sham group did not change across testing (see Figure 7B).

Path circuity of the left- and right-hands during the withdraw phase of movement were measured across testing (see Table 5). A Repeated-measures ANOVA conducted on left-hand path circuity failed to reveal any significant difference. However, changes were observed in right-hand path circuity during the withdraw phase of movement.

A Repeated-measures ANOVA conducted on right-hand path circuity revealed a significant main effect of Day [F (2, 16) = 4.814, p = 0.023, η^2^p = 0.376] yet failed to reveal a significant main effect of Group or Group by Day interaction. Post hoc trend analysis revealed that path circuity increased from day 3 to day 7 and decreased from day 7 to day 14 for both groups [*F* (1, 8) =5.763, *p* = 0.043, η^2^p =0.419].

The concentration of movement with the left- and right-hand was measured on day 3, 7, and 14 for both groups (see Table 5). A Repeated-measures ANOVA conducted on left-hand parameter of concentration revealed a significant main effect of Group [F (2, 16) = 7.813, p = 0.023, η^2^p = 0.494] yet failed to reveal a significant main effect of Day or Group by Day interaction. Rats in the sham group demonstrated more concentrated movement with the left-hand during the withdraw phase of movement, while rats in the MCAO group exhibited less concentrated left-hand withdraws across testing. Yet a Repeated measures ANOVA conducted on right-hand concentration reveled similar organization of movement between groups across testing.

The previous analysis revealed group differences in the parameter of concentration with the left-hand during the withdraw phase of movement. Therefore, the heading direction of the left-hand (see Figure 9G) was not statistically evaluated during the withdraw phase due to the methodology of circular statistics. However, the heading direction with the right-hand was still evaluated across testing (see Figure 9H). A Repeated-measures ANOVA conducted on right-hand heading revealed a significant Group by Day interaction [F (2, 16) = 4.299, p = 0.032, η^2^p = 0.350] yet failed to reveal a significant main effect of Group or Day. Right-hand withdraws ended toward the left side of the body for sham rats (see Figure 9C), while rats that received a MCAO withdrew closer to the lateral side of the body (see Figure 9D) on day 3 of testing. However, heading did not differ between groups on days 7 or 14 of testing.

## 4. Discussion

A unilateral MCAO produced bimanual disruptions in rat string-pulling behavior. Impaired string-pulling was characterized by persistent bilateral increases in misses with both hands and posture changes. When the string was missed with the contralateral to stroke right-hand, rats with MCAO often demonstrated an open-handed raking-like motion (i.e., extensor spasticity similar to flexor spasticity in patients after stroke) as if unable to close the hand and continued to cycle through subcomponents (i.e., Pulls and Pushes) of movement, as if the string was grasped. MCAO rats also altered their posture by bending and twisting the trunk of the body to complete subcomponents of string-pulling movement during both contacts and misses. In contrast, sham rats exhibited independent arm movements without whole-body assistance and made few misses. Their hands made elliptical up/down excursions to grasp and pull, and their posture remained with minimal sway. In sum, the bilateral rhythmical string-pulling task revealed both hand and proximodistal movement impairments as a consequence of MCAO stroke.

The MCA is a major artery that extends from the internal carotid artery and supplies blood to lateral portions of the parietal, frontal, and temporal lobes. The MCAO lesion model used in the current study occluded the artery just above the rhinal fissure restricting blood flow and inducing damage to the neocortex. The cortical areas irrigated by the MCA include the sensorimotor regions of the trunk, face, and limbs (Navarro-Orozco et al., 2021). Patients that have experienced a stroke within the MCA often exhibit paralyses in the face, arm, or trunk of one side of the body (Ng et al., 2007). In the present study it was found that ∼16.8-30.95% of the ipsilateral somatosensory cortex was damaged by MCAO occlusion, a lesion similar to that produced in other MCAO studies (for review see Witte et al., 2000). Yet most of the motor cortex was spared in the stroke rats.

String-pulling in sham and MCAO rats was organized differently. When string-pulling, sham rats typically remain upright with their torso extended to engage in multiple bilateral rhythmical reaches, string grasps, and string pulls and pushes to advance it into the apparatus. Occasional mouth pulls were interspersed with hand movements. Thus, somatosensory cortex may be expected to be involved in the maintenance of posture for string-pulling, the rhythmical bilateral movements of advancing the string, and the targeted hand grasps used to purchase and release the string (Blackwell et al., 2018b).

MCAO rats approached and pulled the string in a similar amount of time as sham rats suggesting spared function of motivation and attentional processes required to perform the task. Nevertheless, many other aspects of string-pulling were impaired, especially in the hand/arm contralateral to the stroke. The stroke rats missed grasps of the string more frequently with both hands but were especially impaired with the contralateral hand that often failed to close after grasping. The stroke animals made much more use of mouth grasps to assist in advancing the string. Arm movement impairments featured further distances traveled, changes in concentration, and heading direction of movement endpoints. Lifts and advances of the arms were no longer independent but were assisted with whole-body movements, including standing upright extended on the toes and up/down movements of the trunk as well as twisting the whole-body to the left and right while moving downward to advance the string.

Whereas many studies featuring single handed reaching for food by rodents have been used to investigate the consequences of stroke (Schaar et al., 2010; Whishaw et al., 1992; Whishaw et al., 2002; Balbinot et al., 2018), the string-pulling task has multiple positive features as a test for neural contributions to fine motor control. The task requires minimal preliminary training, as rats readily pull on a string placed in their cage as part of normal investigatory behavior. The task provides many iterations of whole-body, arm, and hand movements concurrently. Finally, as shown here, the task can still be performed, albeit with changes, after stroke that compromises nearly all somatosensory cortex. In addition, successful grasps with the hands by MCAO rats were characterized by subcomponents of movement that incorporated the use of the entire body. While these changes were apparent during both left- and right-hand engagement by MCAO rats, they were most apparent when the right-hand attempted to reach, grasp, and withdraw the string. This may indicate a greater limited range in movement on the impaired side. Similar behavioral deficits have been reported in rodent models of MCAO, especially in skilled reaching tasks that assesses each hand, as previously described. These studies have provided translational value for the use of behavioral tasks that involve skilled reaching movements, including bimanual string-pulling behavior. Separate recent studies with rodents and humans suggest that string-pulling behavior is organized similarly into reaches and withdraws to grasp and pull in a string with hand-over-hand motions (Blackwell et al., 2018c; Blackwell et al., 2021; Singh et al., 2019). Thus, this dynamic task affords the opportunity to investigate the effects of stroke and recovery of function in a way that may be more translational to behaviors that both rodents and patients engage in.

Compensation following brain damage is critical for recovery of function. The ability of the brain to compensate following MCAO has been widely reported in the literature with varying levels of success depending on the timing and assessments used (Jones, 2017; Ruan, Yao, 2020). For example, compensation in digit use (Gharbawie et al., 2006) and grasping function (Gharbawie et al., 2008) have been previously observed in skilled reaching tasks that involve the use of one hand following MCAO. Further, skilled reaching tasks typically require extensive training, compared to string-pulling behavior that is acquired within a few days while still providing many repetitions of the behavior (i.e., multiple reaches and withdraws per pulling bout). As such, it may be possible that spontaneous recovery of function or compensation may also occur more quickly and with less training in string-pulling behavior relative to skilled reaching. Since most skilled reaching tasks only assess one hand at a time, there is likely a limit to the development of compensatory responses available following brain damage that support functional recovery. However, the spontaneity and bimanual coordination of the hands and use of the mouth during string-pulling behavior likely recruits more neural structures and results in a greater ability to compensate following damage.

Performance in the string-pulling task revealed both persistent and transient deficits following MCAO providing evidence for potential compensation and functional recovery. Impairments that persisted at 3, 7, and 14 days after MCAO included misses with both hands, left-hand withdraw endpoint concentration, and subcomponents of movement without the string. Although enduring deficits were observed, evidence of compensation was still present. Rats that received a MCAO exhibited greater misses with both hands than sham rats. However, during misses, rats engaged in more advances and lifts with the left-hand which suggests more attempts and effort by the nonimpaired side of the body to initiate engagement with the string. Further, when using the right-hand, MCAO rats often engaged simultaneously in mouth contacts and pulled the string down to the left side once it was in the mouth rather than the right side.

In conclusion, the string-pulling task provides a sensitive assessment to detect changes in fine motor control following MCAO. Several disruptions in string-pulling behavior were observed in these rats. First, the string was missed more often with both hands after MCAO, and when the string was missed with the right-hand, rats continued to cycle through subcomponents (i.e., Pushes) of string-pulling behavior as if the string were grasped in the hand. Second, when the string was missed, these rats also failed to make a grasping motion with the right-hand and instead, demonstrated an open-handed raking-like motion. Third, postural changes were observed after MCAO, such that rats used their entire body suggesting compensatory adjustments to advance the string into the apparatus. Lastly, changes in distance and direction that are critical to movement organization were also observed after MCAO.

Despite these impairments in fine motor skills, motivation to engage in and to complete the string-pulling task to obtain a reward was preserved following MCAO. This work demonstrates the importance of using detailed functional analyses of movement to detect changes in performance. Further, this study provides a foundation for future work to investigate other stroke models and to evaluate the efficacy of therapeutic interventions that have the potential to enhance neuroplasticity.

## Statements and Declarations

The authors do not have any funds, conflict of interests or competing interests to disclose. The datasets generated during the current study are available upon request.

## References

Alaverdashvili M, Whishaw IQ (2008). Motor cortex stroke impairs individual digit movement in skilled reaching by the rat. Eur J Neurosci., 28(2):311–22. https://doi.org/10.1111/j.1460-9568.2008.06315.x.

Balbinot G, Schuch CP, Jeffers MS, McDonald MW, Livingston-Thomas JM, Corbett D (2018). Post-stroke kinematic analysis in rats reveals similar reaching abnormalities as humans. Sci Rep, 8, 8738. https://doi.org/10.1038/s41598-018-27101-0.

Batschelet E. Circular Statistics in Biology, ACADEMIC Press, 111 FIFTH AVE., NEW YORK, NY 10003, 1981 (1981, 388).

Bederson JB, Pitts LH, Tsuji M, Nishimura MC, Davis RL, Bartkowski H (1986). Rat middle cerebral artery occlusion: evaluation of the model and development of a neurologic examination. Stroke, 17(3): 472–476. https://doi.org/10.1161/01.str.17.3.472.

Blackwell AA, Köppen JR, Whishaw IQ, & Wallace DG (2018a). String-pulling for food by the rat: Assessment of movement, topography and kinematics of a bilaterally skilled forelimb act. Learning and Motivation, 61: 63–73. https://doi.org/10.1016/j.lmot.2017.03.010

Blackwell AA, Widick WL, Cheatwood JL, Whishaw IQ, Wallace DG (2018b). Unilateral Forelimb Sensorimotor Cortex Devascularization Disrupts the Topographic and Kinematic Characteristics of Hand Movements While String-pulling for Food in the Rat. Behavioural Brain Research. 338, 88–100. https://doi.org/10.1016/j.bbr.2017.10.014.

Blackwell AA, Banovetz MT, Qandeel, Whishaw IQ, Wallace DG (2018c). The structure of arm and hand movements in a spontaneous and food rewarded on-line string-pulling task by the mouse. Behavioural Brain Research. 345, 49–58. https://doi.org/10.1016/j.bbr.2018.02.025.

Blackwell AA, Schell BD, Osterlund Oltmanns JR, Whishaw IQ, Ton ST, Adamczyk NS, Kartje GL, Britten RA, Wallace DG (2021). Skilled movement and posture deficits in rat string-pulling behavior following low dose space radiation (28Si) exposure. Behavioural Brain Research. 400, 113030. https://doi.org/10.1016/j.bbr.2020.113010.

Cheatwood JL, Burnet D, Butteiger DN, Banz WJ. (2011). Soy protein diet increases skilled forelimb reaching function after stroke in rats. Behav Brain Res., 216(2):681–4. https://doi.org/10.1016/j.bbr.2010.09.013.

Cramer SC, Nelles G, Benson RR, Kaplan JD, Parker RA, Kwong KK, et al. (1997). A functional mri study of subjects recovered from hemiparetic stroke. Stroke, 28, 2518–2527. https://doi.org/10.1161/01.str.28.12.2518.

Edwards LL, King EM, Buetefisch CM, Borich MR (2019). Putting the “Sensory” Into Sensorimotor Control: The Role of Sensorimotor Integration in Goal-Directed Hand Movements After Stroke. Frontiers in integrative neuroscience, 13, 16. https://doi.org/10.3389/fnint.2019.00016.

El Amki M, Baumgartner P, Bracko O, Luft AR, Wegener S (2017). Task-Specific Motor Rehabilitation Therapy After Stroke Improves Performance in a Different Motor Task: Translational Evidence. Translational stroke research, 8(4), 347–350. https://doi.org/10.1007/s12975-016-0519-x.

Gharbawie OA, Auer RN, Whishaw IQ (2006). Subcortical middle cerebral artery ischemia abolishes the digit flexion and closing used for grasping in rat skilled reaching. Neuroscience, 137(4):1107–1118. https://doi.org/10.1016/j.neuroscience.2005.10.043.

Gharbawie OA, Williams Ptja, Kolb B, Whishaw IQ (2008). Transient middle cerebral artery occlusion disrupts the forelimb movement representations of rat motor cortex. European Journal of Neuroscience, 28(5), 951–963. https://doi.org/10.1111/j.1460-9568.2008.06399.x.

Gonzalez CL, Kolb B. (2003). A comparison of different models of stroke on behaviour and brain morphology. Eur J Neurosci., 18(7):1950–62. https://doi.org/10.1046/j.1460-9568.2003.02928.x.

Gonzalez CL, Gharbawie OA, Williams PT, Kleim JA, Kolb B, Whishaw IQ (2004). Evidence for bilateral control of skilled movements: ipsilateral skilled forelimb reaching deficits and functional recovery in rats follow motor cortex and lateral frontal cortex lesions. Eur J Neurosci., 20(12):3442–52. https://doi.org/10.1111/j.1460-9568.2004.03751.x.

Jones TA (2017). Motor compensation and its effects on neural reorganization after stroke. Nature Reviews Neuroscience, 18(5):267–280. https://doi.org/10.1038/nrn.2017.26.

Kleim JA, Boychuk JA, Adkins DL (2007). Rat models of upper extremity impairment in stroke. ILAR J., 48(4):374–84. https://doi.org/10.1093/ilar.48.4.374.

Lai CH, Sung WH, Chiang SL, Lu LH, Lin CH, Tung YC, Lin CH (2019). Bimanual coordination deficits in hands following stroke and their relationship with motor and functional performance. J Neuroeng Rehabil., 16(1):101. https://doi.org/10.1186/s12984-019-0570-4.

Ma R, Xie Q, Li Y, Chen Z, Ren M, Chen H, Li H, Li J, Wang J (2020). Animal models of cerebral ischemia: A review. Biomedicine & Pharmacotherapy, 131. https://doi.org/10.1016/j.biopha.2020.110686.

Navarro-Orozco D, Sánchez-Manso JC. Neuroanatomy, Middle Cerebral Artery. Treasure Island, StatPearls Publishing, 2021.

Ng YS, Stein J, Ning M, Black-Schaffer RM. (2007). Comparison of clinical characteristics and functional outcomes of ischemic stroke in different vascular territories. Stroke, 38(8):2309–14. https://doi.org/10.1161/STROKEAHA.106.475483.

Peterson CCH (2019). Sensorimotor processing in the rodent barrel cortex. Nat Rev Neurosci., 20, 533–546. https://doi.org/10.1038/s41583-019-0200-y.

Ruan J, Yao Y (2020). Behavioral tests in rodent models of stroke. Brain Hemorrhages, 1:171-184. NIHMSID: NIHMS1657874. https://doi.org/10.1016/j.hest.2020.09.001.

Sainburg R, Good D, Przybyla A (2013). Bilateral synergy: a framework for post-stroke rehabilitation. J Neurol Transl Neurosci, 1:1025.

Schaar KL, Brenneman MM, Savitz SI (2010). Functional assessments in the rodent stroke model. Exp & Trans Stroke Med, 2, 13. https://doi.org/10.1186/2040-7378-2-13.

Singh S, Mandziak A, Barr K, Blackwell AA, Mohajerani MH, Wallace DG, Whishaw IQ (2019). Human string-pulling with and without a string: movement, sensory control, and memory. Experimental Brain Research.237, 3431–3447. https://doi.org/10.1007/s00221-019-05684-y.

Trueman RC, Diaz C, Farr TD, Harrison DJ, Fuller A, Tokarczuk PF, Stewart AJ, Paisey SJ, Dunnett SB (2017). Systematic and detailed analysis of behavioural tests in the rat middle cerebral artery occlusion model of stroke: Tests for long-term assessment. J Cereb Blood Flow, 37(4):1349–1361. https://doi.org/10.1177/0271678X16654921.

Whishaw IQ, Pellis SM, Gorny BP (1992). Skilled reaching in rats and humans: parallel development of homology. Behavioral Brain Research, 47: 59–70. https://doi.org/10.1016/s0166-4328(05)80252-9.

Whishaw IQ, Suchowersky O, Davis L, Sarna J, Metz G, Pellis S (2002). Impairment of pronation, supination, and body co-ordination in reach-to-grasp tasks in human Parkinson’s disease (PD) reveals homology to deficits in animal models. Behav Brain Res, 133:165–176. https://doi.org/10.1016/s0166-4328(01)00479-x.

Witte OW, Bidmon HJ, Schiene K, Redecker C, Hagemann G (2000). Review Functional differentiation of multiple perilesional zones after focal cerebral ischemia. J Cereb Blood Flow Metab., 20(8):1149–65. https://doi.org/0.1097/00004647-200008000-00001.

